# Cerebrospinal fluid-driven ependymal motile cilia defects are implicated in multiple sclerosis pathophysiology

**DOI:** 10.1101/2025.07.06.663378

**Authors:** Maxime Bigotte, Adam M.R. Groh, Elia Afanasiev, Vincent Wong, Kevin Lancon, Moein Yaqubi, Finn Creeggan, Craig S. Moore, Adil Harroud, Raphael Schneider, Philippe Séguéla, Marc Charabati, Fiona Tea, Antoine P. Fournier, Yu Chang Wang, Jiannis Ragoussis, Alexandre Prat, Simon Thebault, Stephanie Zandee, Jo Anne Stratton

## Abstract

**Background:** Multiple sclerosis is a neurodegenerative autoimmune disorder of the central nervous system (CNS) in which autoreactive immune cells migrate through a damaged blood brain barrier, resulting in focal demyelinating lesions of both the white and grey matter. Of increasing interest is the repeated observation that beyond focal lesions, there are also diffuse, surface-in gradients of pathology in MS, wherein damage is most severe directly adjacent to cerebrospinal fluid (CSF)-contacting surfaces, such as the subpial and periventricular areas. This observation suggests that toxic factors within MS CSF may be contributing to the emergence and/or evolution of surface-in gradients. Directly separating the CSF from the periventricular parenchyma are ependymal cells – a glial epithelium that are equipped with tufts of motile cilia which are critical for circulating CSF solutes and regulating local fluid flow. While damage to ependymal cilia has the potential to drastically modify CSF homeostasis and thus contribute to the damage of CSF exposed regions, these motile cellular structures have yet to be investigated in the context of MS.

**Methods:** We first conducted single cell RNA sequencing of fresh human periventricular brain tissue containing ependymal cells from MS patients and non-MS disease controls. We subsequently collected CSF from MS patients and exposed cultured rodent ependymal cells to this CSF in order to evaluate impact on ependymal ciliary function. To complement our direct evaluation of cilia in the context of MS, we also confirmed whether cilia were altered in a classic animal model of MS, experimental autoimmune encephalomyelitis (EAE), and also designed a novel transgenic animal model to evaluate the cellular and behavioural effect(s) of adult ependymal ciliary disruption.

**Results:** Single cell RNA sequencing analysis of human ependymal cells in MS demonstrated largescale dysregulation of ciliary genes and *in situ* stains of MS brain tissue confirmed a loss of ependymal cilia. Exposure of ependymal cells to MS CSF led to transcriptional modification of ciliary gene and protein expression and reduced ciliary beating frequency. Likewise, analysis of ependymal cells in EAE also demonstrated altered cilia gene and protein expression. Conditional knockout in adult mice, of the critical cilia-associated gene *Ccdc39* in ependymal cells led to transient ventricular enlargement, increased periventricular microglial density, and alterations in nesting behaviour.

**Conclusion:** These data suggest that motile cilia in ependymal cells are dysregulated in CNS autoimmunity. More importantly, however, they provide evidence to suggest that ependymal cilia disruption could play an active role in the development of periventricular pathology in MS and can lead to behavioural deficits that may underlie aspects non-motor MS symptomatology.

## Introduction

The brain ventricles are lined with multi-ciliated ependymal cells, which have around 50 multiple motile cilia attached to the apex of the cell.^1^ The cerebrospinal fluid (CSF) that circulates within the brain ventricles has protective hydromechanical properties, controls brain interstitial homeostasis, circulates chemicals that control the development and activity of the brain, and contains circulating immune cells.^2^ CSF is mainly produced by the choroid plexuses, but a sizeable proportion is thought to come from interstitial fluid exchange across the ependyma.^3^ A transcriptomic analysis across age, region and species highlighted that cilia motility is likely one of the most preserved functions of ependymal cells.^4^ Motile cilia beat in a coordinated manner and are polarized at the cell level (rotational polarity) and at the tissue level (translational polarity) to produce local flow of CSF.^5^ Numerous studies have shown that developmental blocking of ciliary beating induces ventricular enlargement and hydrocephalus, suggesting an important role of ciliary beating in controlling the volume of the ventricles in development^6–18^. But as of today, ciliary beating is considered only a partial contributor to the bulk flow of CSF in adults compared to cardiac pulsation, respiration and bodily movement. ^19–22^ Rather, it is thought that ependymal cell motile cilia control the microcirculation of solute cues at the surface of the ventricles which forms a compartmentalized functional micro-environment.^23^ This compartmentalized functional micro-environment is speculated to directly impact parenchymal brain cells in the periventricular region, for example modulating the migration of neuroblasts to the olfactory bulb and impacting the activity of neuronal and astroglial networks.^24,25^ In summary, ciliary beating controls major homeostatic brain functions, but their role in most adult disease contexts is not well defined.^1^

Multiple sclerosis (MS) is an autoimmune neuroinflammatory disease of the CNS characterized by the dissemination in time and space of perivenular demyelinating lesions containing autoreactive B-cells, T-cells and macrophages.^26^ Curiously, periventricular lesions appearing around the brains lateral ventricles are a common feature of MS. ^27^ Recent studies have also highlighted the presence of a gradient of pathology appearing in the gray matter around the CSF-brain borders, namely the ependyma and the glial limitans superficialis,^28^ as disease progresses.^29,30^ Indeed, there is a periventricular *surface-in* gradient of pathology in which neuronal and axonal loss, demyelination and recruitment of activated microglia is most pronounced closest to the ventricular surface/ependyma. These findings suggest that toxic factors contained in the CSF, such as immune cells, autoantibodies, pro-inflammatory cytokines and/or thrombotic proteins, may contribute to the formation of periventricular pathology. ^31–33^ Interestingly, work in mice and humans has begun to show that ependymal cell reactivity and associated cell adhesion/permeability changes may be implicated in the pathogenesis of *surface-in* gradients in MS. In autoimmune experimental encephalomyelitis (EAE), the most common preclinical model of MS, reactive ependymagliosis was observed concomitantly with subventricular stem cell alterations, barrier protein expression changes and basal body polarity alterations.^34,35^ In humans, post-mortem tissue analysis of MS patients has suggested that ependymal cells become flattened or are absent.^36^ Recent work from our group confirmed that ependymal cells are morphologically altered in MS, but also demonstrated that periventricular *surface-in* gradients of pathology directly associate with stretches of reactive ependymagliosis that have decreased expression of pan-cadherin, a critical junctional protein.^37^ Multiomic single nucleus RNA sequencing of human ependymal cells in MS revealed widespread junctional protein gene dysregulation driven by discrete transcription factor hubs. These data provide the first evidence that alterations to ependymal cell barrier properties may participate in periventricular pathology formation and/or progression in MS. That said, the role of motile cilia alterations remains to be evaluated.

The objective of the present study is to evaluate if ependymal motile cilia are altered in MS and in MS models and to determine how cilia alterations could be involved in periventricular pathology formation. We demonstrate that ependymal motile cilia are modified in MS and in a preclinical model of disease. Furthermore, we show evidence, in a targeted transgenic mouse model, that adult-induced ependymal motile cilia disruption can lead to periventricular tissue and behavioral alterations with shared features of MS periventricular inflammation and MS symptomatology.

## Materials and methods

### Human tissue samples and processing

For donor information, see Supplementary Table 1. Tissue samples were collected from 4 MS patients and 4 controls (CTRL; 3 amyotrophic lateral sclerosis (ALS) and 1 hereditary spastic paraplegia (HSP) patients) under the medical assistance in dying (MAID) program of Quebec at the Centre de Recherche du Centre Hospitalier de l’Université de Montréal (CR-CHUM). MS diagnosis was determined in accordance with the revised 2017 McDonald’s criteria.^26^ ALS diagnoses was based on the revised El Escorial criteria and postmortem neuropathological assessment.^38,39^ Ventricular/subventricular tissue was dissected from the frontal horn of the lateral ventricle from coronal sections human brain autopsy by or under direct supervision of certified pathologists and neurologists. Part of the tissue was freshly dissociated for single cell RNA extraction and sequencing, and another part was fixed in 10% formalin and kept at 4°C for 1-7 days until paraffin embedding. Additional formalin-fixed tissue from the Douglas-Bell Canada Bain Bank (DBCBB) was used for histology analysis (4 MS patients and 4 non-inflammatory disease patients). In both cases, approval of patient’s brains for use in research was obtained prior to death, and tissue was processed according to the ethical approvement of the local committee (Research Ethics Boards of the Montreal Neurological Institute-Hospital and the McGill University Health Centre (protocol approval number (BH07.001 and 2023–9159). Formalin-fixed tissues were dehydrated using a standard set of alcohol and xylene washes and then embedded in paraffin. Cassettes containing the paraffin-embedded brain tissue were stored at room temperature until cutting for immunostaining. 5uM thick sections were prepared using a microtome.

### Tissue dissociation, RNA extraction and library preparation for single cell RNA sequencing from human periventricular area

The ventricular/subventricular dissected samples were immediately weighed, minced using scalpels, dissociated into single cell suspension using the Adult Brain Dissociation Kit (Miltenyi Biotec) according to the manufacturer’s protocol but without the red blood cell removal step. The isolated single cells were then counted using a hemocytometer and assessed via LIVE/DEAD viability testing (Thermo Fisher).

The single cells were processed, and single-cell libraries were generated using the 10x Genomics Chromium Controller and Chromium Next GEM Single Cell 5’ Reagent Kits (10x Genomics), as per the manufacturer’s protocol. The sequencing-ready libraries were quality controlled for size distribution and yield (LabChip GX Perkin Elmer) and quantified using qPCR (KAPA Biosystems Library Quantification Kit for Illumina platforms). Libraries were loaded on NovaSeq 6000 and sequenced using the following parameters: 100 bp Read1, 10 bp Index i7, 10 bp Index i5 and 100 bp Read2.

### Single cell RNA sequencing bioinformatics

Demultiplexing, genome alignment, and gene quantification were performed with Cell Ranger ARC 50. Specifically, data was processed using the cellranger count pipeline and aligned against the human genome (GRCh38) to generate count matrices.^39^ The resulted count matrices were subjected to subsequent data processing, normalization, batch integration, and gene expression in Seurat (v.5.0.1)^40^ in R (v.4.1.2). Briefly, raw counts were normalized with the LogNormalize method (scaling factor = 10,000), and the top 2,000 highly variable genes were identified using a variance-stabilizing transformation (VST). Data were scaled, and principal component analysis (PCA) was performed using these variable features. The first 20 principal components were used to construct a k-nearest neighbor (KNN) graph, followed by clustering with the Louvain algorithm (resolution = 0.4). Uniform Manifold Approximation and Projection (UMAP) was applied for dimensionality reduction and cluster visualization, with clusters displayed based on their seurat_clusters identity. Differentially expressed genes (DEGs) between clusters were calculated using the Wilcox test implemented in Seurat. DEGs were selected using an adjusted *P*-value of less than 0.05 and a log2 fold-change threshold of 0.25. Gene Ontology (GO) analysis of DEGs was performed using clusterProfiler.^41^

### Cilia-related GO term analysis

Following determination of differentially expressed genes for a dataset, genes were categorized as either “ciliary” or not by assessing if they resided with the GO term gene set for “cilium organization” (GO:0044782). Following this, GO term over-representation analysis within the biological process (BP) category was run separately for significantly (adjusted *P*-value < 0.05) up- and down-regulated genes. GO terms were filtered as “cilia” relevant if they were considered offspring of the GO term for “cilium organization” (GO:0044782).

### Patient CSF

MS patient CSF was collected in 3 different locations and were used as 3 different cohorts. CSF was quickly put on ice after the spinal tap and centrifuged at 400g at 4°C for 10min. After spin, the cell pellet was frozen, and the CSF supernatant was aliquoted and stored at −80°C right after collection and was thawed only once prior usage. CSF collected from patients with non-inflammatory neurological disease (NIND) was used as controls. Patient’s consent was collected prior use of specimens for research and The Code of Ethics of the World Medical Association (Declaration of Helsinki) was followed. Cohort 1 contained 4 MS CSFs and 2 NIND CSFs provded by Dr. Craig S. Moore (Memorial University of Newfoundland) and were used for bulk RNA sequencing of ependymal cells after treatment with CSF *in vitro*. Cohort 2 contained 5 MS CSF and 2 NIND CSF and was sourced at the Clinical Biospecimen Imaging and Genetic Repository (CBIG) at the Montreal Neurological Institute (MNI, McGill University) and was used for immunolabeling analysis after treatment on ependymal cells *in vitro*. Cohort 3 contained 7 MS CSF and 3 NIND CSF and was sourced at St. Micheal’s Hospital of Toronto and graciously sent by Dr. Raphael Schneider (St. Micheal’s Hospital, Toronto) and used for live ciliary recording of ependymal cells after treatment with CSF *in vitro*. CSF donor information is present in the Supplementary Table 2.

### Primary culture of rat ependymal cells and treatments

Rat primary cultures of ependymal cells were obtained as previously described.^40,41^ Cells were considered mature after 21 days *in vitro*. Then patient CSF was applied during 48 hours at a 1:1 dilution ratio before collection for bulk RNA sequencing, fixation for immunolabeling or live cilia video recordings. Non-treated (NT) cells, or cells treated with artificial CSF (aCSF) containing 124 mM NaCl, 2 mM KCl, 26 mM NaHCO3, 1.8 mM MgSO4, 1.25 mM NaH2PO4, 10 mM Glucose, 1.6 mM CaCl2, pH 7.4, were also used as controls.

### RNA isolation and bulk RNA sequencing of ependymal cells from primary cultures

Cells were washed with Hank’s Balanced Salted Solution (HBSS) and buffer RL was then added to lysate the cells. Cells lysates were stored at −80°C until further steps. RNA isolation was then performed using the Norgen Singel Cell RNA Isolation Kit (Norgen 51800). After isolation, RNA concentration was assessed using a NanoDrop spectrophotometer (ThermoFischer Scientific), and samples were diluted to 50 ng/µL and kept at −80°C.

All library preparation and next generation sequencing was performed by Genome Quebec. Libraries were constructed in a stranded fashion, with additional PolyA enrichment. Illumina Novaseq paired-end sequencing with 100 pb read length was requested, with 50M reads per sample. FASTQ output files were then copied onto Compute Canada for long term storage. Transcript counts were generated using the salmon aligner, with GRCr8 assembly used as reference.^42^ Salmon outputs were read in using the tximeta R library and differential gene expression analysis was conducted using the DESeq2 R library.^43,44^ Cell ranger outputs were downloaded from GEO accession number GSE199460 and read in using the Seurat library.^45–48^ Following correction of mislabeled cell identifiers in the provided metadata (as described previously in),^35^ counts were normalized using a Wilcoxon signed-rank test as implemented in the Seurat FindMarkers function. Functional term over-representation analysis was conducted using the clusterProfiler R library, with GSEA calculations conducted using the fgsea R library.^49^

### Live cilia recording of ependymal cells from primary cultures

Ependymal cells from primary cultures were placed under an epifluorescent microscope (Axio Observer, Zeiss) equipped with a high-speed camera (Fastcam mini AX50, Photron) under a 40x objectives and bright light. Videos were recorded using the PFV4 software (version 4.0.6.2, Photron) at 500 frame/s. Videos were saved as .avi.

### Animals

All animal work received prior approval (protocol #2019-8102) and was carried out in accordance with the guidelines and regulations of the local Animal Care Committee. Animals were ordered from Charles River. All animals were kept on a 12-hour light/dark cycle and had access to food and water *ad libitum*. Mice were randomized, no animals were excluded, and sample size calculations were not performed. A total of 51 animals were used. 8 C57BL/6 mice were used to induce MOG-EAE (and no immunization controls), 12 C57BL/6 mice were used for intracerebroventricular injections, 15 double transgenic *αSMACreER^T^*^2^*::ROSA26^tdTomato^* mice and 13 triple transgenic *αSMACreER^T^*^2^*::ROSA26^tdTomato^*::*Ccdc39^flox/flox^*mice were used for *Ccdc39* knock-out experiments and 4 Sprague Dawley pregnant rats with 8-12 pups were used for primary rat ependymal cell cultures.

### Experimental autoimmune encephalomyelitis

C57BL/6 mice were immunized against MOG_35-55_ as previously described.^35^ MOG_35-55_ peptide powder (MOG35-55-P-10; Alpha Diagnostic) was prepared at 4 mg/mL (14025076; Gibco). Incomplete Freund’s adjuvant (DF0639-60-6; BD Difco) was mixed with desiccated mycobacterium (DF3114-33-8; BD Difco) and diluted to 8 mg/mL. MOG and incomplete Freund’s adjuvant were mixed for 2 h and then left at 4°C overnight to create complete Freund’s adjuvant. On day 0, emulsions were injected subcutaneously with 50 µL of emulsion into each hind flank (100 µL total per animal) of C57BL/6 mice (*n* = 11; female). A working solution of pertussis toxin (PHZ1174; Invitrogen) was made by dilution in HBSS (14025076; Gibco) at 2 µg/mL. On day 2, 200 µL of pertussis toxin working solution was injected intraperitoneally into each animal. MOG-EAE mice were killed at 11–17 days post-immunization (peak disease, *n* = 11 females) by cardiac perfusion with PBS while anesthetized with 1%–3% isoflurane. Littermate controls were not given injections (*n* = 8; females).

### EAE single cell RNA sequencing dataset analysis

Single cell RNAseq outputs were downloaded from GEO accession number GSE254863^35^ and read in using the Seurat R library.^45–48^ Cells annotated as ependymal cells in the provided annotations were subsetted, and the resulting dataset was down sampled to contain an equal number of cells from each source (500). Counts for this subsampled dataset were then normalized and scaled as described previously,^50^ and differentially expressed genes between the EAE and control conditions were determined using the FindMarkers function exposed by the Seurat R library.

### Transgenic *Ccdc39* knockout mouse model

*Ccdc39* is a gene required for assembly of the inner dynein arm and dynein regulatory complex in all motile cilia, including those expressed by ependymal cells. To induce knockout of *Ccdc39* specifically in ependymal cells, we crossed *Ccdc39^flox/flox^* mice and *αSMAC-reER^T^*^2^*::ROSA26^tdTomato^* mice, which were sourced from Dr. June Goto (University of Cincinnati) and Dr. Jeff Biernaskie (University of Calgary), respectively. Triple transgenic animals homozygous for *Ccdc39*, *αSMACre*, and *tdTomato* were utilized for evaluation of the down-stream effects of adult ependymal Ccdc39 knockout on brain health. Adult CreER^T2^ activity was induced by five consecutive intraperitoneal injections (24 hrs apart; 1mg/injection) of tamoxifen (Z)-4-Hydroxytamoxifen; H7904, Sigma-Aldrich). A total of 28 animals in 4 cohorts were used for the study. Cohort 1 was composed of 3 WT animals (2F;1M) and 3 *Ccdc39*^flox/flox^ animals (3M) sacrificed 2 weeks after tamoxifen injection. Cohort 2 was composed of 4 WT animals (3F;1M) and 3 *Ccdc39*^flox/flox^ animals (3M) sacrificed 4 weeks after tamoxifen injection. Cohort 3 was composed of 3 WT animals (2M;1F) and 4 *Ccdc39*^flox/flox^ animals (4F) sacrificed 4 weeks after tamoxifen injection. And cohort 4 was composed of 5 WT animals (5M) and 3 *Ccdc39*^flox/flox^ animals (3F) sacrificed 8 weeks after tamoxifen injection.

### Mouse behavioral testing

Nesting behavior was evaluated as described previously.^51^ In short, mice were transferred to individual cages without environmental enrichment and a single 3g pressed cotton nestlet 1h before the dark phase. The nests were then scored the next morning using a scale of 1-5, with 0 representing no nest and 5 representing a characteristic nest.

Novel object recognition was evaluated using a three-day protocol adapted from Leger & colleagues.^52^ Before beginning the protocol, animals were individually placed into the testing apparatus (45cm x 45cm x 50cm box/open field) for 5 minutes per day for three days to acclimatize. The apparatus was cleaned between each mouse using a mixture of 50% ethanol in water. This cleaning was repeated between animals for the entirety of the protocol (both testing apparatus and object). On day 1 of the protocol, mice were habituated once more in the testing apparatus without objects for 10 minutes. On day 2, two identical objects were placed in the center of the testing apparatus and mice were individually placed in the apparatus facing away from the objects and left to explore for 5 minutes. On day 3, one of the identical objects was replaced with a novel object and mice were once again individually placed in the testing apparatus and left to explore. Mouse activity within the testing apparatus on days 2 and 3 was recorded using a webcam so that time spent interacting with each object could be evaluated. The preference index was calculated using the following equation: time with novel object / (time with familiar object + time with novel object).

Y-maze assessment was achieved by acclimatizing the animals to the y-maze and testing room 7 days prior to testing, and again on the testing day itself, 1h prior to test start time, as described previously.^53^ On testing day, the arms of the y-maze were labeled A, B, and C, and each mouse was placed in the distal end of arm A facing the center of the maze to start the test. The mice were then left to freely explore the maze for 8 minutes. The maze was thoroughly cleaned with 50% ethanol in water between each test. Percent alternation was calculated using the following formula: (number of alternations / [total number of arm entries – 2]) x 100.

### Immunolabeling and histology

Human tissue immunolabeling was performed on MAID tissue as previously described,^37^ with mouse anti-FoxJ1 (1:500; 14-9965-82; eBioscience) primary antibodies. For cilia morphology, the cytoskeletal Bielschowsky staining was performed on DBCBB tissue, as previously described.^37^ Mouse tissue fixation, cryo-sectioning and immunolabeling were performed as previously described,^35,37^ with rabbit anti FoxJ1 (1:200; ab235445; abcam) or rabbit anti Iba1 (1:1000; 019-19741; Wako) primary antibodies. After 48h treatment, rat primary cultures of ependymal cells were fixed and stained as previously described,^35,54^ with mouse anti-FoxJ1 (1:500; 14-9965-82; eBioscience) primary antibodies.

### Microscopy and image analyses

Immunolabeled mice brain slices were imaged with an epifluorescent microscope (AxioObserver, Zeiss) and images were analyzed using Fiji (National Institute of Health). Immunolabeled primary cultured cells were imaged using a ImageXpress Mini confocal microscope (Molecular Devices). Images were then analyzed using the MetaXpress software (version 6.2.0.66, Molecular Devices). Ciliary beating frequency was measured using Fiji by building kymographs via the Reslice [/] function. Each ciliary beating cycle corresponds to one pic on the kymograph and the ciliary beating frequency (CBF) was calculated by measuring the distance between two pics (*l*) using the following formula: CBF = 1/ (*l* x δ*t*) (CBF: beating/s; *l*: pixel length between two pics and δ*t*: 1/camera’s frame rate).

### Statistical analysis

Graphs and statistical tests were performed with Graphpad Prism (version 10.4.0). Normality was tested using Shapiro-Wilk tests prior to choosing between parametric and non-parametric tests. According to the result, t-tests or Mann-Whitney tests were used to compare groups of two, and ordinary one-way ANOVA or Kruskal-Wallis tests were used to compare multiple groups. Post-hoc multiple group tests were performed as well (Fisher’s LSD test for ANOVA or Dunn’s for Kruskal-Wallis). Graphs are plotted as means + 95% confidence interval (CI) histograms superposed to individual data points. *P*-values < 0.05 were considered statistically significant and the level of significance is indicated as follows: *=*P*≤0.05; **=*P*≤0.01; ***=*P* ≤0.001 and ****=*P*≤0.001.

### Data availability

The code used to conduct all bioinformatic analyses is available on demand. MOG-EAE and human MS single cell RNA sequencing data sets are publicly available under the GEO accession number GSE254863 and GSE301585 and at https://singlocell.openscience.mcgill.ca.

## Results

### Postmortem single cell RNA sequencing and *in situ* analysis of ependymal cells reveals cilia alterations in multiple sclerosis

To investigate if ependymal motile cilia are altered in MS, we performed single cell RNA sequencing from ventricular/subventricular samples freshly resected from the dorsal horn of the lateral ventricles of freshly autopsied brains of MS patients (*n* = 4) and non-MS patients as controls (*n* = 4) (Fig. 1A). For donor information, see Supplementary Table 1. Ependymal cells were found in each patient and were one of 6 captured cell populations, identified by expression of *FoxJ1* and *PIFO* (Fig. 1B, C, D). We performed differential gene expression analysis across all MS versus control ependymal cells and found that within all DEGs, 125 cilia related transcripts were significantly downregulated and 23 were upregulated in MS compared to controls (Fig. 1E). Most of the top 50 cilia DEGs were associated with axonemal and dynein arm proteins such as *Ccdc39*, which we later used to build our transgenic model (Fig. 1F). 9 Cilia GO-terms were significantly downregulated, and 1 cilia GO-term was significantly upregulated in MS compared to controls (Fig. 1G). To confirm that there was evidence of cilia modifications at the protein level, we performed histological stains on periventricular MS tissue. We found a reduction in FoxJ1 protein expression (Fig. 1H) and a significant reduction in the number of cilia at the apex of ependymal cells using a Bielschowsky stain (Fig. 1I and J, *P* = 0.0107).^37^ Altogether, this work suggests that motile cilia in ependymal cells are modified in MS at the transcriptomic, protein and morphological level.

**Figure 1.**
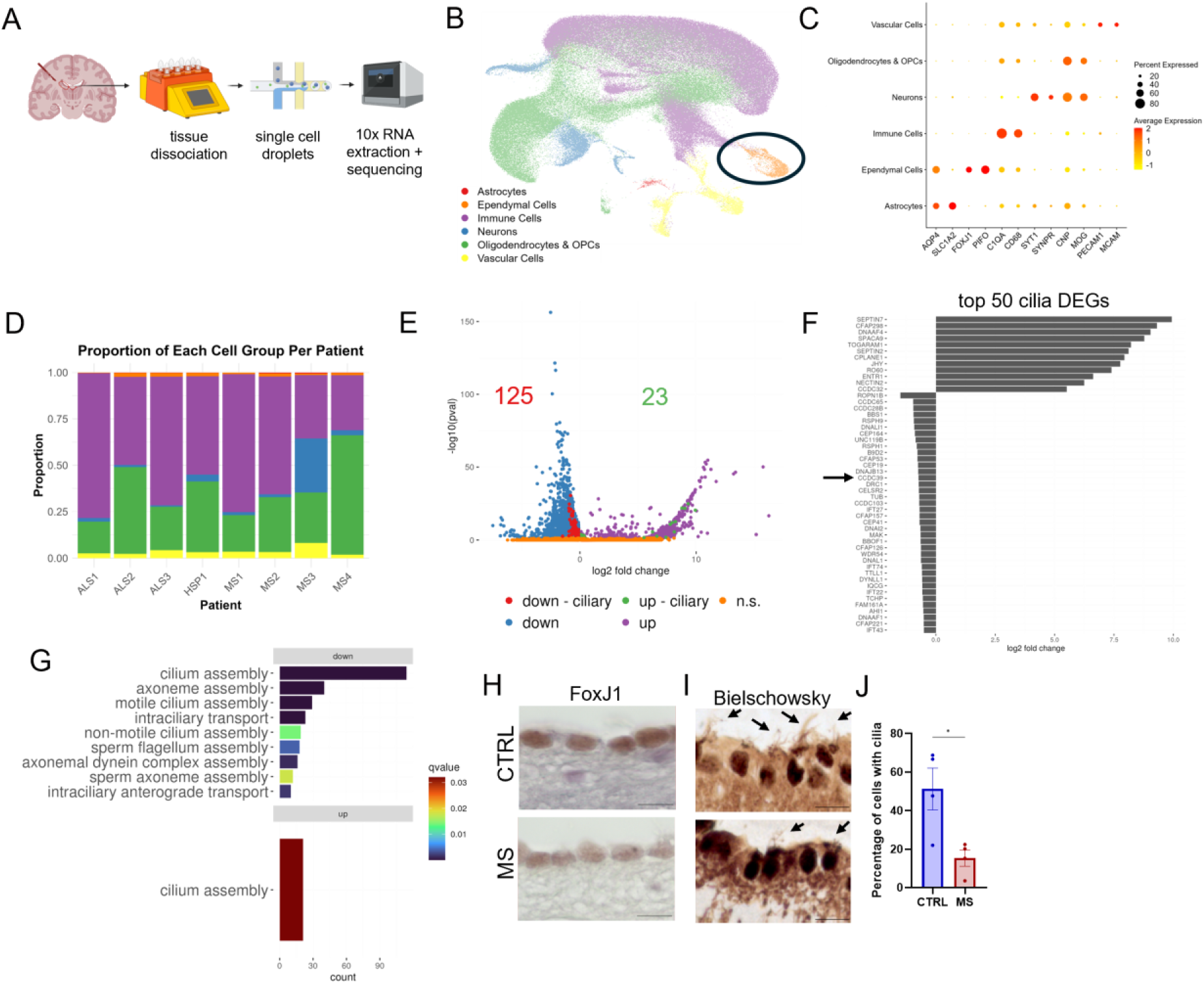
Transcriptomic and histological analysis of motile cilia of ependymal cells from MS postmortem tissue. (**A**) Schematic workflow of tissue dissociation and sequencing of dissected ventricular/periventricular pieces from MS patients (*n* = 4) and CTRL patients after rapid autopsy (*n* = 4). (**B**) UMAP plot of single-cell transcriptomic profiles where ependymal cells were identified (black circle) based on *FoxJ1* and *PIFO* expression. (**C**) Dot plot of periventricular cell type-associated gene expression. Dot colour indicates the average expression value, whereas dot size indicates the percentage of cells expressing the gene. (**D**) Stacked bar plot showing normalized proportions of cell types as defined in panel B per patient. (**E**) Volcano-plot of the total DEGs and specific motile cilia DEGs in ependymal cells in MS compared to NIND (threshold: *Log2FC* <-0.25; *q*-value < 0.05). (**F**) Bar plot of the top 50 most significant motile cilia DEGs in ependymal cells in MS compared to CTRL. (**G**) Bar plot of the number of significant upregulated and downregulated GO-terms generated from the list of significant motile cilia DEGs using clusterProfiler. (**H**) Microscopic micrographs of ependymal cells from the lateral ventricle of MS patients stained for FoxJ1 (scale bar = 10 µm). (**I**) Representative microscopic micrographs of ependymal cells from the lateral ventricle of MS patients compared to CTRL patients where motile cilia were identified by a Bielschowsky staining (black arrow, scale bar = 10 µm). (**J**) Histogram of the mean ± 95% CI of the percentage of cells with cilia quantified in Bielschowsky stained slides.

### MS CSF disrupts ependymal cilia *in vitro*

MS CSF is hypothesized to drive pathology in CSF-exposed brain regions, driving surface-in damage and ependymal cell disruption.^29,37^ Therefore, to determine whether the transcriptomic, protein and morphological changes observed in MS were also associated with a functional disruption in cilia motility, we applied CSF from MS patients on rat primary cultures of ependymal cells at a 1:1 dilution for 48 hours and compared their effects to non-treated cells, artificial CSF and CTRL CSF (Fig. 2A). Bulk RNA sequencing after 48 hours exposure also showed that, within all total DEGs, 47 cilia related genes were significantly downregulated, and 71 cilia related genes were significantly upregulated (Fig. 2B). Interestingly, important ciliogenesis transcription factors and elements of the axoneme were transcriptionally modified (Fig. 2C). Cilia GO-term analysis showed that the most modified pathways were associated with cilium assembly, and that cilia-associated pathways were only upregulated and not downregulated (Fig. 2D). Noably, after 48 hours exposure with MS CSF, FoxJ1 expression was significantly increased compared to controls (Fig. 2E and F, *P* = 0.0083), yet there was reduced ciliary beating frequency after 48 hours compared to controls (Fig. 2G, *P* = 0.0333). These results demonstrate that MS CSF contains elements able to rapidly induce motile cilia gene/protein and functional alterations in ependymal cells.

**Figure 2.**
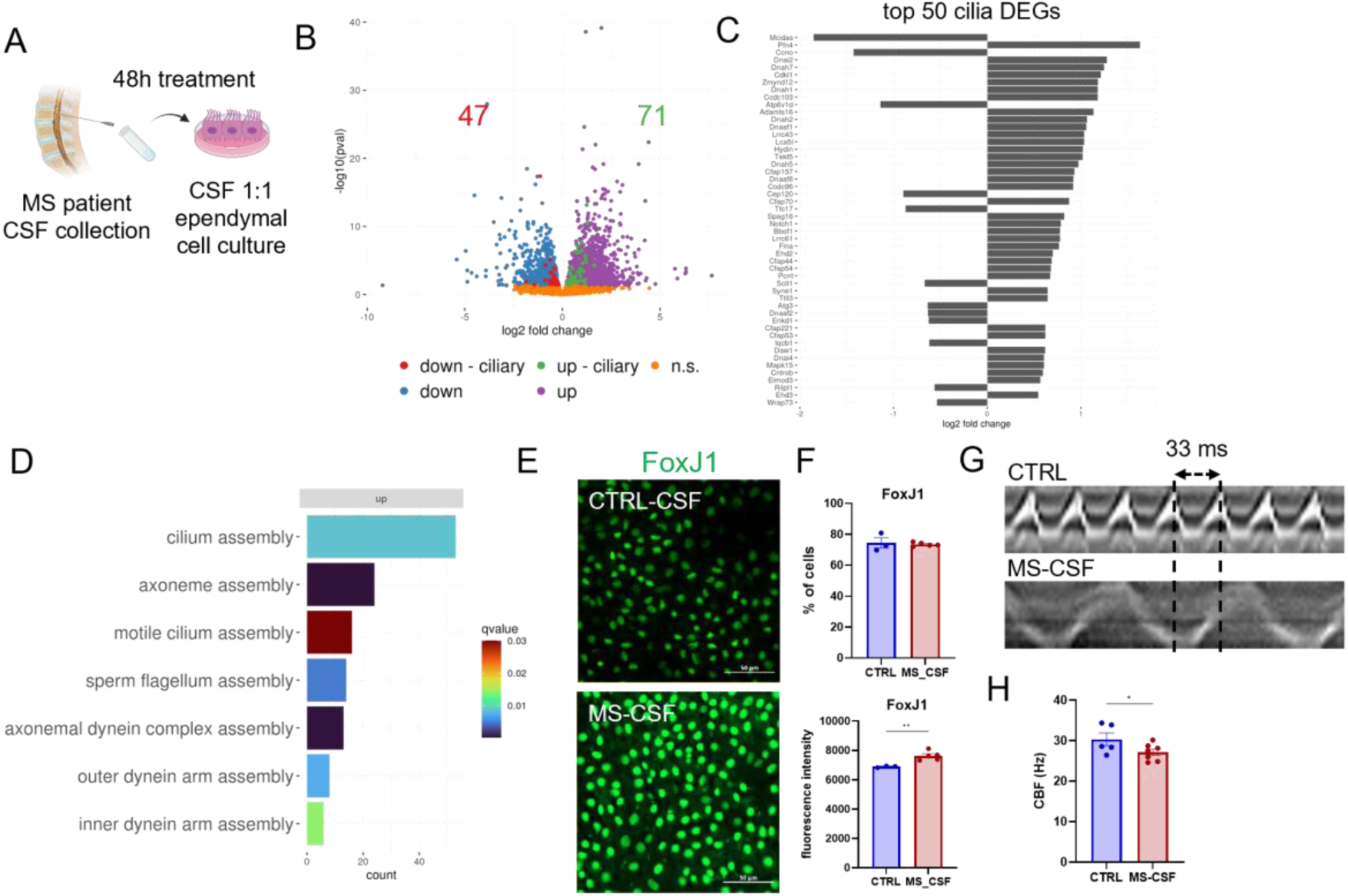
Analysis of motile cilia genes, protein and beating frequency after MS CSF treatment on rat primary cultures of ependymal cells. (**A**) Illustration of CSF collection and treatment 1:1 on rat primary ependymal cell cultures for 48h. (**B**) Volcano-plot of the total DEGs and specific motile cilia DEGs obtained by RNA bulk sequencing of ependymal cells after 48h treatment with MS CSF in comparison to non-treated cells (threshold: *Log2FC* <-0.25; *q*-value < 0.05). (**C**) Bar plot of the top 50 most significant motile cilia DEGs in ependymal cells after 48h treatment with MS CSF compared to non-treated cells. (**D**) Bar plot of the number of significant upregulated GO-terms generated from the list of significant motile cilia DEGs using clusterProfiler. (**E**) Representative microscope micrographs of rat primary ependymal cell cultures stained for FoxJ1 (green) after 48h treatment with MS-CSF or CTRL-CSF. (**F**) Histograms ± 95% CI of the mean percentage of FoxJ1 expressing cells and the mean fluorescence intensity of FoxJ1. Each data point is the mean of triplicate measurements done in different culture wells. (**G**) Representative kymographs (200 ms) of one motile cilia tuft from rat primary ependymal cell cultures after 48h in a control condition or treated with MS-CSF. (**H**) Histogram ± 95% CI of the mean beating frequency (*Hz*) of motile cilia of primary ependymal cell cultures 48h after treatment with MS-CSF compared to control conditions. Each data point is the mean beating frequency of 10 different motile cilia measured in triplicates in 3 different culture wells.

### Transcriptomic and in situ analyses confirm that cilia are impaired in an animal model of MS

To further investigate motile cilia in central nervous system autoimmunity, we used a whole brain single cell RNA sequencing dataset generated from MOG_35-55_-EAE animals and matched controls previously explored by our group (Fig. 3A and B).^35^ After subsetting the ependymal cell cluster and performing DEG analysis between EAE and controls, we found 8 cilia related genes were significantly downregulated and 53 cilia related genes were significantly upregulated (Fig. 3C). Of these genes, 47 genes with the 50 highest fold change were in fact upregulated and included several dynein arm and axonemal proteins (Fig. 3D), including *FoxJ1* (*Log2FC* = −0.79, *P*adj = 4.69E.38) and *Ccdc39* (*Log2FC* = 0.65, *P*adj = 1.13E-10). Of the top cilia DEGs, most were associated with of cilia-associated GO-terms such as cilium assembly and intraciliary transport, and that cilia-associated pathways were only upregulated and not downregulated (Fig. 3E). Further, FoxJ1 protein expression was downregulated in ependymal cells of MOG_35-55_-EAE animals at peak disease scores compared to controls (Fig. 3F and G, *P* = 0.0005). Like previously found,^55–58^ brain lateral ventricles were enlarged at disease peak (Fig. 3H and I, *P* = 0.032). Interestingly, we also noticed that lateral, 3^rd^ and 4^th^ ventricle enlargement was more pronounced in the caudal part of the brain compared to the rostral lateral ventricles (Supplementary figure 1). These results show that transcriptional and protein expression modification to ependymal cilia are also present in an animal model of MS and that these changes are associated with ventricular enlargement, a feature of MS.

**Figure 3.**
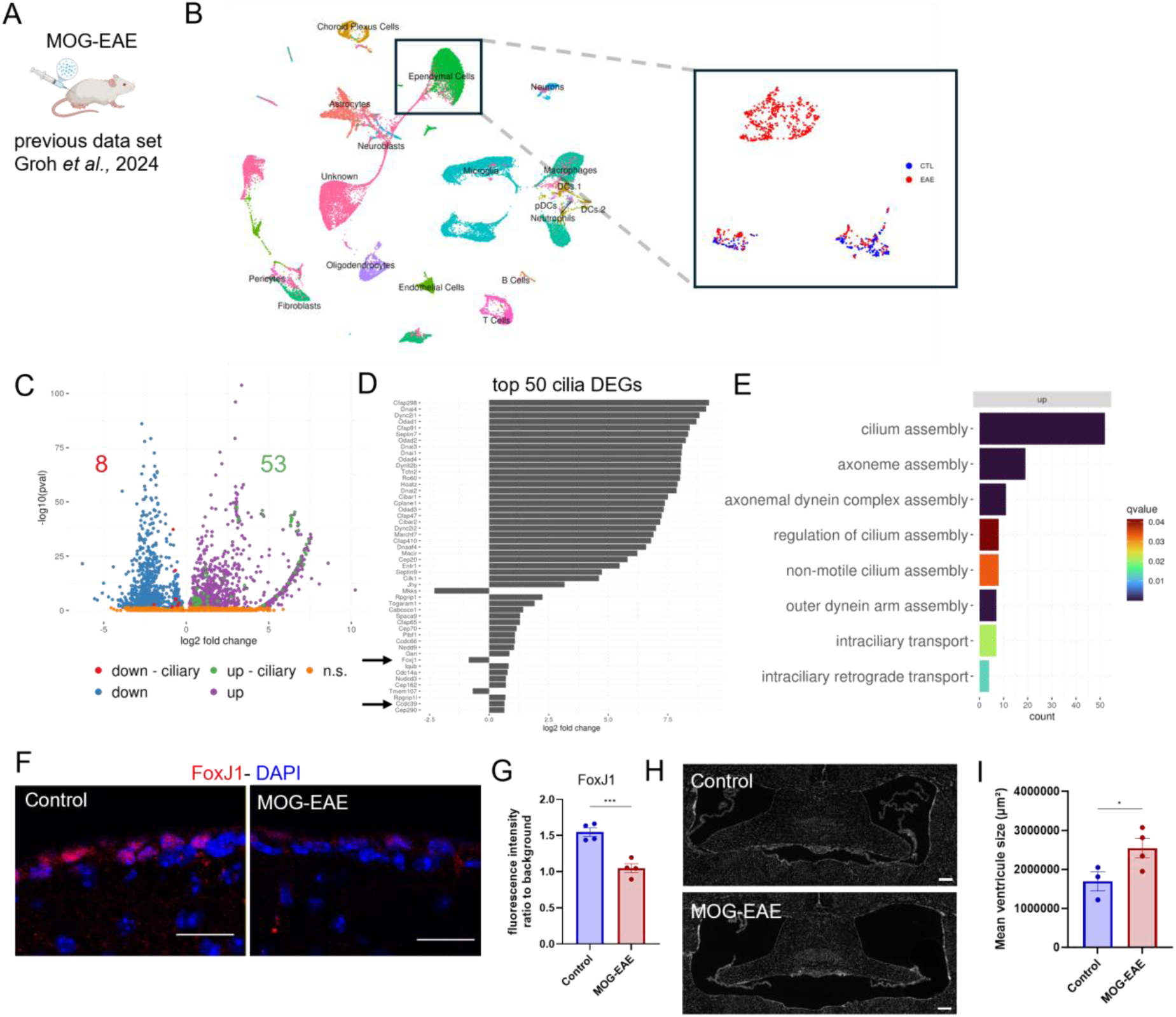
Single cell RNA sequencing and tissue analysis of motile cilia of ependymal cells in MOG_35-55_-EAE animals. (**A**) Single cell RNA sequencing data of ependymal cells from MOG_35-55_-EAE animals from Groh *et al.,* 2024 was reanalyzed for motile cilia specific genes. (**B**) Two UMAPs were generated to plot all the cells present in the data set and ependymal cells were clustered using *FoxJ1*. The first UMAP represents all the cell clusters and the second UMAP represents ependymal cell clusters from control and MOG_35-55_-EAE animals. (**C**) Volcano-plot of the total DEGs and specific motile cilia DEGs within the ependymal cell cluster of the single cell RNA sequencing dataset of MOG_35-55_-EAE compared to control animals (threshold: *Log2FC* <-0.25; *q*-value < 0.05). (**D**) Bar plot of the top 50 most significant motile cilia DEGs in ependymal cells of MOG_35-55_-EAE compared to control animals. (**E**) Bar plot of the number of upregulated significant GO-terms generated from the list of significant motile cilia DEGs using clusterProfiler. (**F**) Representative microscopic microphotographs of ependymal cells labeled with FoxJ1 (red) from the lateral ventricle of MOG_35-55_-EAE and control animals (scale bar = 20 µm). (**G**) Histogram ± 95% CI of the mean FoxJ1 fluorescence intensity. Each data point is the mean fluorescence per animal normalized to background intensity. (**H**) Representative microscopic mosaic microphotographs of the whole anterior lateral ventricle + dorsal 3^rd^ ventricle (between −0.2 to −0.5 mm to Bregma in the antero-posterior axe) of MOG_35-55_-EAE and control animals labeled with DAPI (gray, scale bar = 200 µm). (**I**) Histogram ± 95% CI of the mean ventricular area (µm²) of MOG_35-55_-EAE and control animals. Each data point is the ventricular area of one animal.

### Inducible knock-out of *Ccdc39* prompts periventricular alterations and cognitive deficits in mice

To test if motile cilia dysfunction in ependymal cells alone could lead to pathological changes relevant to MS, we designed a transgenic mouse model in which *Ccdc39* was knocked out specifically in ependymal cells by tamoxifen injection in *Ccdc39^flox/flox^*-*αSMAC-reER^T^*^2^*::ROSA26^tdTomato^* mice. *Ccdc39* is coiled-coil domain-containing protein involved in the assembly of the inner dynein arm which has been previously associated with postnatal cilia defects and hydrocephalus in knock-out mice,^6^ and was significantly downregulated in ependymal cells from both MS patients (Fig. 1D, black arrow) and MOG-EAE mice (Fig. 3D). We observed ventricular enlargement at 2 weeks and 4 weeks after tamoxifen injection in transgenic animals compared to oil injected transgenic mice (Fig. 4A and B, 2 weeks *P* = 0.003, 4 weeks *P* =0.012 and 8 weeks *P* = 0.333). We also investigated if adult-onset *Ccdc39* knock out could lead to periventricular alterations, a characteristic feature of MS pathophysiology. In MS, a key periventricular pathological feature is the presence of microglia, which is observed in periventricular regions even beyond *surface-in* gradients.^35,37^ Interestingly, we observed a signification increase of periventricular Iba^+^ microglial cells (Fig. 4D and E, 2 weeks *P* = 0.005, 4 weeks *P* = 0.008 and 8 weeks *P* = 0.283).

**Figure 4.**
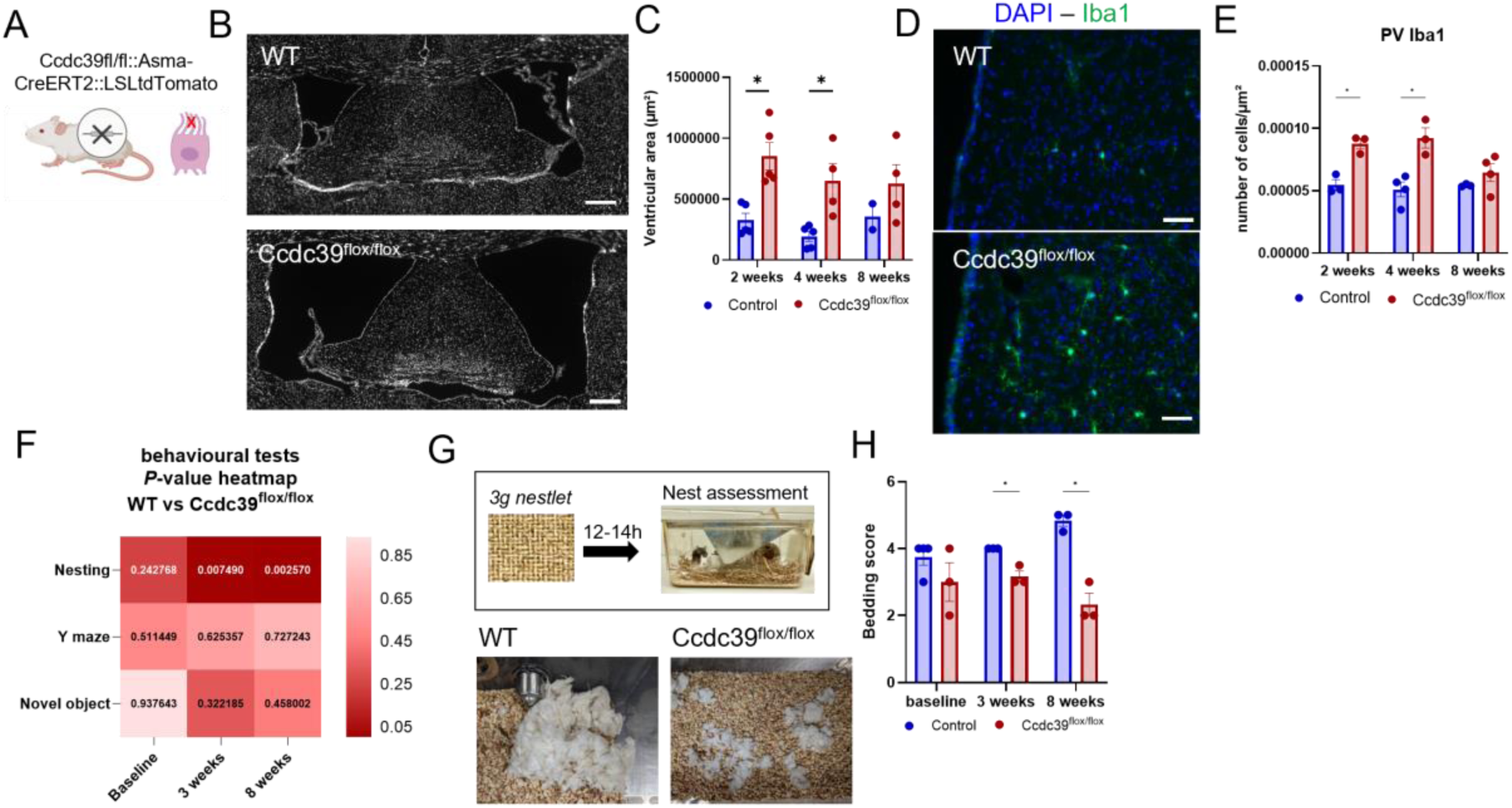
Tissue and behavioral analysis of mice that underwent a *Ccdc39* conditional knock out. (**A**) Schematic showing that double transgenic *αSMACreER^T^*^2^*::ROSA26^tdTomato^* mice were crossed with *Ccdc39^flox/flox^* to specifically induce *Ccdc39* knock-out by tamoxifen injection in ependymal cells. (**B**) Representative microscopic mosaic microphotographs of the whole anterior lateral ventricle + dorsal 3^rd^ ventricle (between −0.2 to −0.5 mm to Bregma in the antero-posterior axe) of *Ccdc39^flox/flox^* animals compared to WT (gray = DAPI, scale bar = 250 µm). (**C**) Histogram ± 95% CI of the mean ventricular area (µm²) of *Ccdc39^flox/flox^* animals compared to WT at 2, 4 and 8 weeks after tamoxifen injections. Each data point is the ventricular area of one animal. (**D**) Representative microscope photographs of *Ccdc39^flox/flox^* and WT periventricular Iba1 positive cells (green) (scale bar = 50 µm). (**E**) Histogram ± 95% CI of the mean number of cells/µm² of periventricular Iba1 positive cells at 2-, 4- and 8-weeks post tamoxifen injection. Each data point is the number of cells/µm² for one animal. (**F**) Heatmap of the T-test *P*-values for each behavior test scores of *Ccdc39^flox/flox^* compared to WT animals at baseline, 3 weeks and 8 weeks post tamoxifen injection. (**G**) Schematic of the nesting behavior test and representative photographs of nests at 12-14h after starting the nesting test for *Ccdc39^flox/flox^* and WT animals. (**H**) Histogram ± 95% CI of the mean bedding score (size of nests) at baseline, 3 and 8 weeks after tamoxifen injection. Each data point is the mean bedding score for each animal.

MS patients experience a gamut of non-motor symptoms such as fatigue, depression and cognition alterations that are not well-understood.^59^ Alterations to CSF dynamics and subsequent ventricular enlargement are a candidate explanation for some of this symptomatology.^60–62^ To evaluate this possibility, we also tested if ependymal-specific *Ccdc39* knockout leads to cognitive impairment by assessing animal behavior with Y-maze, novel object recognition and nesting behavioral tests (Fig. 4F). Interestingly, mice showed a significant reduction in nesting scores at 3 weeks and 8 weeks after tamoxifen injection (Fig. 4F-H, baseline *P* = 0.326, 3 weeks *P* = 0.038 and 8 weeks *P* = 0.007). Altogether, these results showed that alteration of ciliary beating of ependymal cells in adult mice was sufficient to produce ventricular enlargement, an increase in periventricular microglia numbers and to induce non-motor behavior phenotype changes.

## Discussion

While lateral ventricle-adjacent periventricular lesions have been observed by MRI for decades,^27^ the discovery of periventricular *surface-in* gradients of pathology is recent.^29,30^ Both these observations suggest that toxic elements within the CSF of MS patients could drive the formation of this lesion type; however, how the ependymal brain-CSF border is involved in this process has largely been left unstudied. In a recent study from our group, such *surface-in* gradients of pathology were found to be associated with ependymal cell pathology.^37^ While it was shown that ependymal cells underwent a reactive gliotic phenotype associated with deterioration of adherens junctions, whether cilia motility is also impaired remains unaddressed. In this follow-up study, we provide evidence that ependymal motile cilia are compromised at the transcriptomic and protein level in both MS and EAE, and that MS CSF applied on cultured ependymal cells can drive acute cilia alterations. Furthermore, we show that reducing ependymal cilia motility in adult mice is sufficient to reproduce some MS-like periventricular pathology and behavioral alterations. This suggests that ependymal cilia alterations in MS may not only be a consequence of CSF cytotoxicity but can also participate in periventricular pathogenic mechanisms.

Although we showed that ependymal motile cilia are altered in MS and MOG-EAE, and that MS CSF was able to trigger these alterations, the exact mechanisms leading to cilia defects remain unknown. Cilia alterations could be part of the reactive phenotype that ependymal cells undertake in response to inflammation (i.e., ependymagliosis).^35^ The exact trigger of the ependymal cell reactive phenotype remains to be found but our CSF experiment suggests that CSF can be a source of such a trigger. In our previous study,^37^ Nichenet analysis revealed that proinflammatory cytokines such as interferon-γ (IFNγ) or tumor necrosis factor α (TNFα) could be drivers. This is consistent with work from Magliozzi and colleagues which showed that periventricular *surface-in* gradients are associated with elevation of IFNγ and TNFα in MS CSF, but other factors are also likely playing a role, such as autoantibodies.^63^

Regardless of the initiating trigger, the precise temporal interplay between ependymagliosis, barrier protein disruption, and ciliary defects remains to be elucidated—whether these alterations arise concurrently or unfold as a sequential cascade of events. If we suppose that inflammatory factors are directly causing acute ependymal cell reactivity, it is logical that ependymagliosis may happen first, followed by barrier breakdown and cilia alterations. Indeed, in a forebrain targeted EAE model, ependymal cell ‘swallowing’ and GFAP expression in ependymal cells started at 3 days post induction, followed by junctional protein expression changes and cilia basal body delocalization starting at 7 days post induction.^34^ Reorganization of transcriptional priority in response to inflammation would impact important functions in ependymal cells, thus explaining why barrier proteins and ciliary beating would be affected.^35^ It is possible that this inflammatory environment is leading to de-differentiation of ependymal cells into their precursor state, as radial glial precursor cells, that re-expresses GFAP and re-acquire progenitor properties, such as primary cilium. De-differentiation would be associated with a loss of the maintenance of specialized transcriptional pathways that are critical for ependymal cells, such as cilia motility gene programs.^4^ Primary cilium of radial glia cells usually disappears as they commit to ependymal cell fates, as their basal body divides to give rise to multiple motile cilia when differentiating during development.^64–66^ It is tempting to speculate that this process is reversed in MS - we observed striking upregulation of *several* cilia-related gene programs in our dataset (not just programs for cilia motility) which could represent sensory cilia gene programs related to ependymal cell reverting back to this developmental origin. Consistent with this idea, others have shown that in the absence of FoxJ1 in adult ependymal cells, cells de-differentiate into a radial-glia phenotype.^67^ It is also possible that changes in mechanical load are contributing to such changes. Ependymal cell morphology or “stretching” is observed in MS and EAE,^34,36^ and in other inflammatory contexts,^54^ which can arise due to high pressure environments leading to higher mechanical forces and cytoskeleton remodeling at the cell membrane. This can lead to cell adhesion molecule and gap junction disruptions inducing cilia anchoring defects.^68,69^ Given cilia anchoring and beating is highly dependent on the maintenance of the apical actin cytoskeleton^70^, this hypothesis fits well. That said, our in vitro work (where mechanical load should be constant) also demonstrated changes in cilia motility suggesting an active regulatory role of factors within the MS CSF. Regardless of the precise mechanisms underlying our observations, these alterations seem to be present long term in MS since ependymagliosis, barrier breakdown and cilia alterations are all found in postmortem tissue at chronic stages of disease.^36,37^ More studies are needed to understand the intrinsic mechanisms that are at play in ependymal cells in the context of neuroinflammation.

Ventricular volume increases were observed both in MOG-EAE, as in previous studies,^56–58^ and in *Ccdc39* conditional knockout mice. When interpreting such observations in the context of the changes we observed relating to cilia motility, our data suggest a role for disrupted cilia motility in ventricular volume changes in MS. In MS, ventricular volume increases are consistently observed in all ventricles^71^, but the diversity of mechanisms driving this have not been extensively explored, beyond the general idea that it is a result of brain atrophy. It is likely that volume changes are region and context specific in MS, and our data suggests that such changes could, in part, be due to defects in ependymal cells. While the MS data we present is focused on the frontal horns of the lateral ventricles due to the fact that periventricular lesions are often located around these horns in MS,^72^ periventricular lesions are also commonly found throughout the brain for example around the 3^rd^ ventricles.^29^ It would be of great interest to understand the regional specificity of ciliary beating alterations in MS to better understand the link between differences in regional ventricular volume increases and periventricular lesion formation, as well as how this relates to clinical outcomes. Interestingly, the horns of the lateral ventricles are regions where the CSF circulates slowly,^19^ and this is the same region in the aging brain that shows increased mechanical stress and ependymal thinning, making the horns more susceptible to periventricular white matter pathology.^73,74^ Are these unique features of the horns contributing to lesion formation in these regions? Ependymal cilia also vary in number and size along the whole ventricular system,^75,76^ and specific flow patterns are orchestrated by oriented ciliary beating patterns in the 3^rd^ ventricles of rodents and pigs.^23^ A recent study showed that a high level of microflows were present in the periphery of the lateral ventricle system (i.e., the frontal and posterior horns), while the majority of cilia were aligned with bulk flow in the rest of the lateral ventricles.^77^ Are these unique region-specific features also contributing to outcomes in MS?

CSF clearance is also reduced in MS^78^ which would lead to alteration in bulk flow, but it is unclear if ependymal ciliary beating alteration would suffice. While ciliary beating is probably sufficient to control CSF circulation in foramens where the diameter is small,^22^ the small mechanical forces that cilia produce is probably insufficient to control CSF bulk flow across all ventricles.^19^ Instead, choroid plexuses CSF production and clearance, cardiac pulse, pulmonary respiration and gravity are key contributors to CSF bulk flow.^20,21^ Since choroid plexus inflammation is also present in MS and correlates with periventricular lesions,^79^ it is conceivable that CSF bulk flow alterations in MS could be reduced due to reduced CSF production. That said, we speculate that there might also be a role of ependymal cilia due to alterations in solute exchange between the interstitial fluid and the CSF at the ependyma.

Even if the exact role of ependymal cells in MS is unclear, changes in these cells in MS could alter important brain functions. Indeed, it could explain why we observed nesting impairments in *Ccdc39^flox/flox^*mice, which was also found previously in EAE and is associated with a depressive phenotype.^80^ Mice with hydrocephalus and ependymal cilia defects develop cognitive impairments,^6,81^ and patients with primary cilia dyskinesia develop age-related ventriculomegaly associated with cognitive deficits, attention deficits and depression,^62^ both suggesting a direct link between ependymal motile cilia deficits and neurological/psychiatric alterations.^82^ Since periventricular MS lesions observed by MRI are associated with cognitive deficits and depression in MS,^60,61^ ependymal motile cilia alterations could participate via these other roles to the formation of cognitive deficits, fatigue and depression observed in MS.

This study shows that motile cilia alterations in MS may not be an inflammatory bystander, but rather may actively contribute to pathogenic evolution in MS. A better understanding of how motile cilia contributes to the maintenance of periventricular tissue health and to the normal cognitive functioning of the brain is needed to understand how their alteration could contribute to the development of periventricular lesions and MS symptoms. This could support the discovery of new therapeutic avenues for cilia-targeted therapies, in diseases like MS where motile cilia are disrupted.

## Acknowledgements

The authors thank Christel Dias for collecting patient consent for CSF processing and for collecting patient information. They thank Hong Li for her expertise in human brain tissue processing and sectioning. They also extend gratitude to Dr. June Goto (University of Cincinnati) and Dr. Jeff Biernaskie (University of Calgary) for providing transgenic animals, to the staff at the Center for Neurological Disease Models (CNDM) at The Neuro for taking care of animals prior to surgery, and to Vanessa Omana (lab manager) for her contributions. Many thanks to Bettina Zierfuss, Romain Cayrol and Wendy Klement from the CHUM for their help with tissue collection for single cell RNA seq cell processing. They are also grateful for feedback provided by Drs. G.R. Wayne Moore and Jack P. Antel. Lastly, they offer sincere appreciation to the DBCBB/MAID brain and CSF banks, and to the donors themselves for their valued contribution to science.

## Supplementary material

Supplementary material is available at *Brain* online.

---

**Supplementary table 1.**
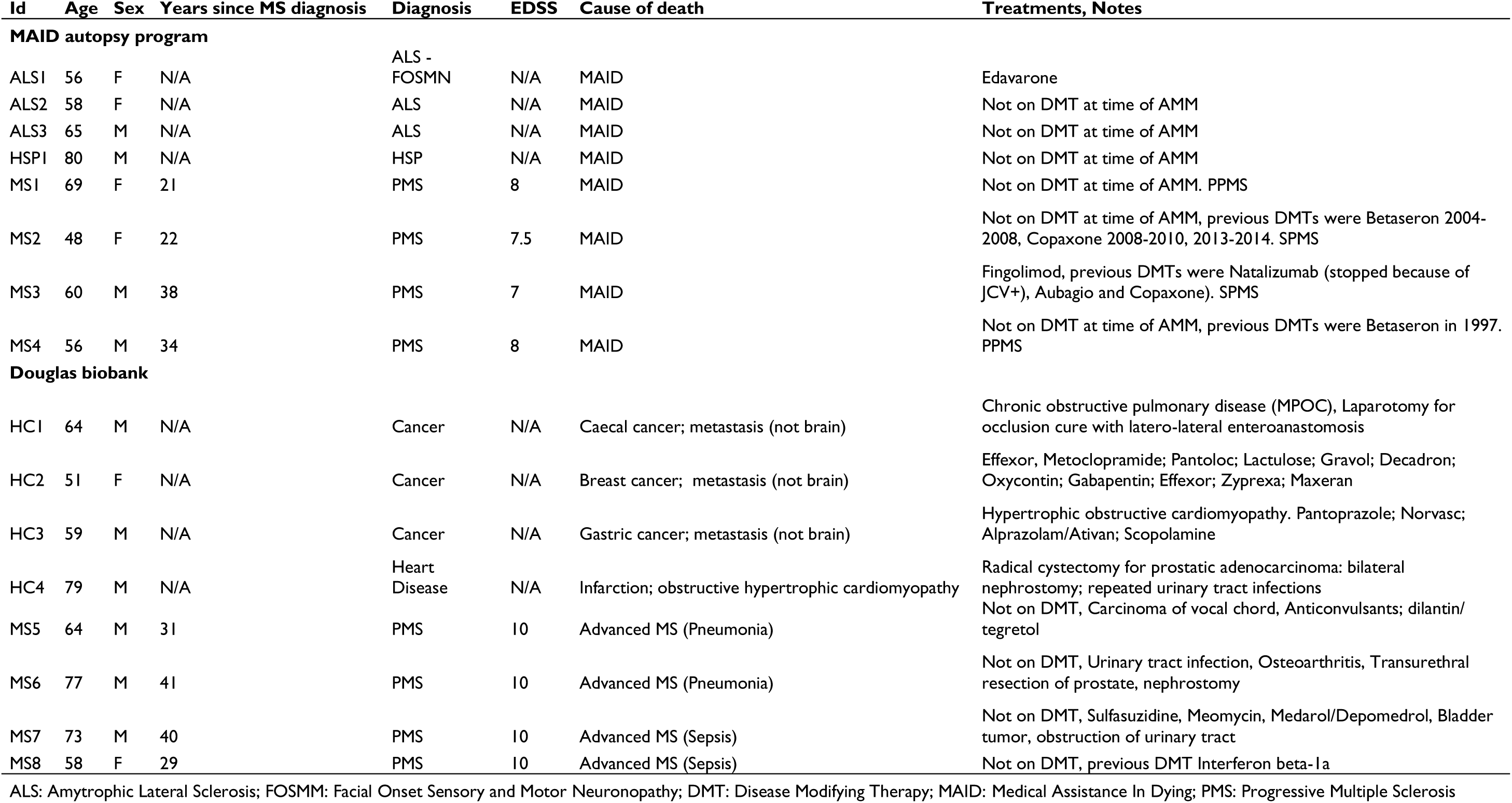
Brain donors information.

**Supplementary table 2.** CSF donors information.

| Age | Sex | Final diagnosis | EDSS | Treatment |
| --- | --- | --- | --- | --- |
| 19 | M | RRMS | 1 | None |
| 45 | F | RRMS | 3 | None |
| 59 | F | RRMS | 3.5 | None |
| 44 | F | RRMS | 1 | Teriflunomide |
| 59 | F | PPMS | 1 | None |
| 43 | F | RRMS | 1 | None |
| 45 | F | RRMS | 2 | None |
| 50 | F | RRMS | 2 | None |
| 37 | M | RRMS | 2 | None |
| 38 | M | RRMS | 2 | None |
| 46 | F | RRMS | 2.5 | None |
| 51 | F | PPMS | 3.5 | None |
| 45 | M | PPMS | 6 | None |
| 30 | M | RRMS | 6.5 | Ocrelizumab |
| 46 | M | PPMS | 7 | None |
| 46 | M | SPMS | 7 | None, Ocrelizumab (discontinued) |
| 50 | F | NIND - neurogenic bladder | N/A | None |
| 37 | F | NIND - vascular | N/A | None |
| 34 | F | Idiopathic intracranial hypertension | N/A | None |
| 18 | F | Migraines+POTS+long COVID | N/A | None |
| 63 | F | non-specific white matter lesion | N/A | None |
| 48 | F | non-specific white matter lesion | N/A | None |
| 45 | F | non-specific white matter lesion | N/A | None |

NIND: Non-Inflammatory Neurological Disease; POTS: Postural Orthostatic Tachycardia Syndrome; PPMS: Primary Progressive Multiple Sclerosis; SPMS: Secondary Progressive Multiple Sclerosis; RRMS: Relapse-Relmitting Multiple Sclerosis

