## Supplementary material for "Cerebrospinal fluid-driven ependymal motile cilia defects are implicated in multiple sclerosis pathophysiology"

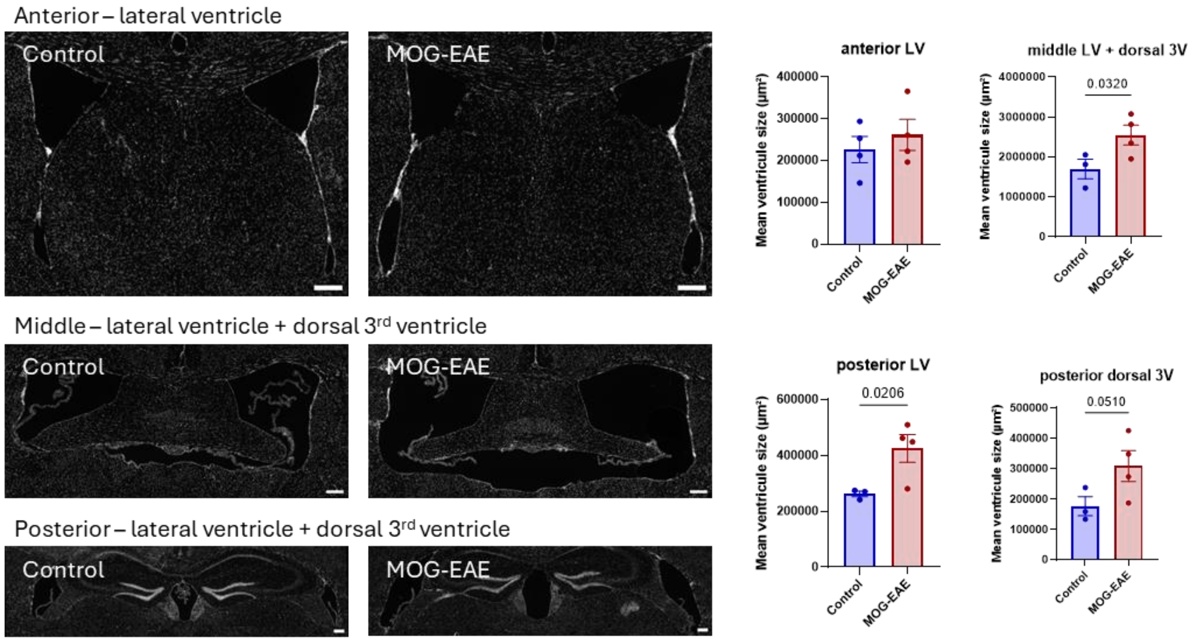


**Supplementary figure 1 Regional ventricular enlargement in MOG-EAE.** Representative microscopic mosaic microphotographs of the anterior lateral ventricle, the anterior + dorsal 3rd ventricle (middle brain), and the posterior lateral and dorsal 3rd ventricle of $MOG_{35\text{-}55}$-EAE and control animals labeled with DAPI (gray, scale bar = 200 µm). Histogram ± 95% CI of the mean ventricular area (µm²) of $MOG_{35\text{-}55}$-EAE and control animals. Each data point is the ventricular area of one animal.
